# RAPID: Real-time image-based autofocus for all wide-field optical microscopy systems

**DOI:** 10.1101/170555

**Authors:** L. Silvestri, M. C. Müllenbroich, I. Costantini, A. P. Di Giovanna, L. Sacconi, F. S. Pavone

## Abstract

Autofocus methods used in biomicroscopy are either based on the search of an optimal focus position – which requires suspending data collection during the optimization process – or on the continuous monitoring of the position of a fiducial plane – which may not coincide with the sample itself. Here, we introduce RAPID (Rapid Autofocus via Pupil-split Image phase Detection), a method for real-time image-based focus stabilization, applicable in all wide-field microscopy systems. We demonstrate that RAPID maintains high image quality in various settings, from *in vivo* fluorescence imaging to light-sheet microscopy. RAPID provides a universal autofocus solution for automated microscopy, and enables quantitative assays otherwise impossible in a standard microscope, such as 3D tracking of fast-moving organisms.

---

Automated operation is increasingly required in optical microscopy systems to expand data and sample throughput. Indeed, image collection is an almost automatic process in many different applications, including whole-slide histological imaging^1^, high-content screening^2^, long-term *in vivo* imaging of cell cultures^3^, high-resolution light-sheet microscopy (LSM)^4^, and tracking assays^5^. In these applications, autofocus (AF) methods are used to maintain image sharpness throughout the acquisition by correcting focal shifts that are introduced by thermal drifts, uneven stages or slides, and by the samples themselves or their motion in a 3D environment.

AF techniques used in microscopy are primarily classified into two categories (Supplementary Fig. 1 and Supplementary Note 1). In image contrast-based methods, the same field of view is imaged at different focal positions, and the position corresponding to the image with the highest contrast is labeled ‘in-focus’^6^. Since a large number of images (typically of the order of 10) have to be collected to establish the best focus, these techniques are not suitable for the correction of fast defocus events (e.g., when tracking living specimens in 3D), and in general significantly reduce system throughput. Conversely, in triangulation methods, the reflection of an oblique light beam from the sample coverslip is used to measure the position of the coverslip itself along the optical axis, which is then used to correct the sample position^7^. Although this approach provides real-time correction, it actually measures the position of the reflective surface and not that of the sample itself; it is therefore impractical when these distances are not fixed (e.g., when the specimens are mounted on gel substrates) or when no reflective surface is present (as in LSM).

The real-time monitoring of the actual focal state of a sample can be achieved by using phase detection, an AF method largely used in photography^8^. The key idea behind this approach is that rays passing through distinct portions of the system pupil intersect the image plane at different lateral positions when the object is defocused (Fig. 1a). By discriminating those two sub-bundles of light rays, the information of their mutual displacement can be used to determine the focal state of the image being acquired by the microscope. Albeit the phase detection principle has been occasionally employed in microscopy, previous implementations were designed around very specific applications, i.e. 3D single-molecule tracking^9^ and whole-slide imaging^10^, preventing a more general use. Indeed, in the former case, the phase detection algorithm starts with the localization of single-molecule centroids, which is clearly inapplicable in other imaging contexts. By contrast, the system described in ref. 10 selects the portions of the pupil by using pinholes, thus preventing its use in low-light settings (Supplementary Note 2).

**Figure 1.**
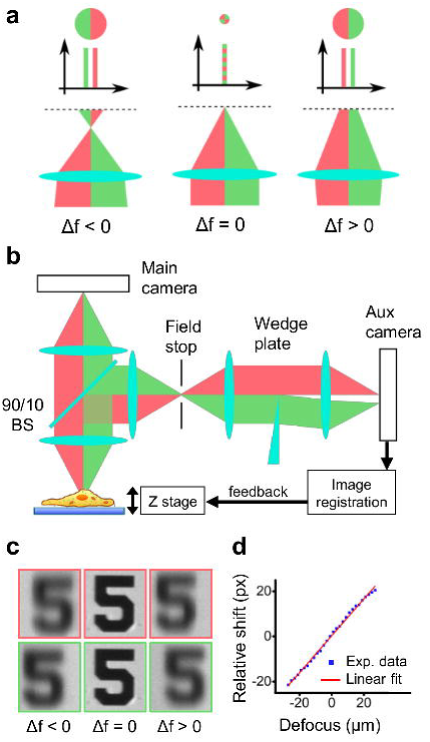
RAPID operation. Lateral motion of the center of mass of rays passing through different pupil portions, depicted in red and green (a). Implementation of RAPID in a standard wide-field microscope (b). Lateral shift of pupil-split images (c). Experimental shift plotted as a function of defocus, together with a linear fit (d).

Here, we introduce RAPID (Rapid Autofocus via Pupil-split Image phase Detection), a phase-detection AF system applicable to all optical microscopy techniques that are based on wide-field detection, including bright-field (BF), epifluorescence (EF), and light-sheet illumination. Differently from other AF methods used in microscopy, this technique provide real-time image-based correction of defocus, and afford reliable operation even at very low light levels, regardless of the kind of samples imaged. We demonstrate RAPID in standard AF applications, as well as in biomicroscopy assays that are beyond the capacity of state-of-the-art methods.

## Results

### RAPID operation

RAPID physically separates rays passing through the two halves of the microscope pupil using a wedge plate, and rigidly aligns the two images produced by the two sub-bundles (Fig. 1b). The quantitative relation between the lateral motion of RAPID images and defocus can be obtained by considering separately the two ray bundles originated from the two halves of the pupil (Supplementary Fig. 2). Depending on the amount of defocus ∆*f*, the lateral displacement of the center of mass of ray bundles (with respect to the central ray) is given by:

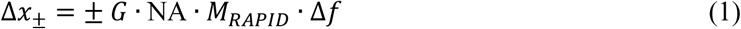

where ± refers to the two different halves, NA is the numerical aperture of the microscope objective used, *M_RAPID_* is the effective magnification in the image space of the RAPID system, and *G* is a geometric factor that is dependent on the shape of the pupil portion used and on the light distribution in the pupil. In the case of two perfect halves of a uniformly filled circular pupil, *G* is given by 4⁄3π (the center of mass of half a unity circle); in general, *G* is of the order of unity. From Eq. (1), it turns out that the mutual distance *d* between the centers of mass of the two ray bundles is linearly dependent on defocus:

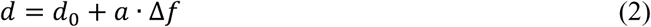

*d*_0_ being the in-focus mutual distance and with *a* = 2 ∙ *G* ∙ NA ∙ *M_RAPID_*.

The above analysis has been derived assuming a single point source. However, since modern microscopes are usually telecentric, such analysis is valid for all the points in the field of view (and in particular *G* is constant throughout the field of view). Therefore, Eq. (2) provides also the mutual displacement of the two bidimensional images produced onto the RAPID auxiliary camera. As far as the system is perfectly telecentric and spatially homogeneous (i.e. neglecting vignetting and distortion), the mutual displacement between the two images is perfectly rigid (Fig. 1c and Supplementary Video 1).

We experimentally verified the relation between *d* and ∆*f*, obtaining *d* by means of cross-correlation (see Methods and Supplementary Methods). Lateral shift between pupil-split images was found to be linearly dependent on defocus in a variety of illumination conditions, objective magnification and NA (Fig. 1d, Supplementary Figs. 3-5). Exploiting this linear behavior, it is possible to infer the focal state of the system from *d* by inverting eq. (2), and use this information to keep the system focused using a simple feedback loop.

The RAPID system can be inserted in any existing microscope by simply adding a beam splitter after the detection objective (Fig. 1b). In this way, using only a small fraction of the detected light (typically 10%, obtained with off-the-shelf beam splitters), RAPID provides instantaneous readouts of the focal state of the system while continuously collecting images with the main camera. Because focus discrimination is reduced to a well-studied computational problem, i.e., image registration, RAPID can capitalize on several established methods^11^ to achieve reliable, flexible, and robust operation (Methods and Supplementary Methods), even far from focus and in low-light regimes. Indeed, we observed reliable focus discrimination over a range 70 times larger than the objective depth of focus and with signal-to-background and signal-to-noise ratios as low as 1.7 and 4.5, respectively (Supplementary Fig. 5 and Supplementary Note 3). The measured focus discrimination accuracy was approximately 70% of the depth of focus and can theoretically be reduced further (Supplementary Note 3 and Supplementary Table 1).

### RAPID autofocusing in whole-slide imaging

To evaluate the flexibility of RAPID, we performed a series of experiments, starting with a standard AF application in optical microscopy: whole-slide histological imaging. In this test, we imaged samples of an atherosclerotic human carotid and human keloid under BF illumination. Unevenness of the mounting slide and XY translation stage can introduce severe defocus when imaging millimeter- or centimeter-sized slides^1^. RAPID provided fast correction of specimen- and system-induced defocus across a large range (several tens of microns), guaranteeing high-contrast imaging in the entire slide without increasing the acquisition time (Fig. 2).

**Figure 2.**
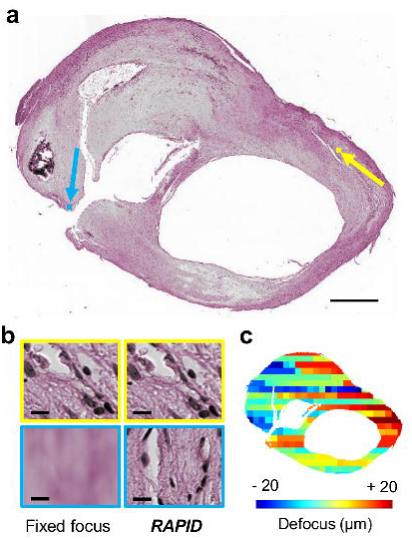
RAPID autofocusing in whole-slide imaging. Application of RAPID in whole-slide histological imaging of an atherosclerotic human carotid (a); insets corresponding to the yellow and blue arrows are shown in (b). Defocus map showing large defocus variability across the slide (c). Scale bars: 1 mm (a), 10 µm (b).

### RAPID focus stabilization in long-term imaging of cell cultures

To examine the stability of RAPID operation over time, we tested the system in another application that commonly employs AF: long-term *in vivo* imaging of a cell culture. We observed a culture of *S. cerevisiae* for more than 12 h in a BF configuration (Fig. 3a, Supplementary Video 2) and confirmed that RAPID maintained image sharpness throughout the imaging session. In contrast, without focus correction, the images became blurred after only 3 h. We repeated the same experiment using a transgenic strain of *S. pombe* expressing tubulin-linked GFP in an EF configuration (Fig. 3b, Supplementary Video 3). Again, RAPID was able to maintain image sharpness for the entire imaging time (> 12 h), in contrast to fixed focus imaging, demonstrating that this method can also perform with the low light fluxes that are typical of *in vivo* fluorescence imaging.

**Figure 3.**
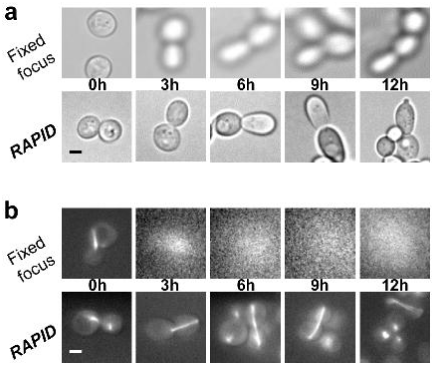
RAPID focus stabilization in long-term imaging of cell cultures. Images of cultured yeast cells taken at different time points show long-term focus stabilization with RAPID in bright-field (a) and epifluorescence (b) microscopy. Scale bars: 1 µm.

### RAPID allows 3D tracking of fast-moving specimens

In the above standard AF settings, the triangulation or image contrast AF methods could also have been used with comparable performances (Supplementary Table 2). Therefore, we evaluated RAPID in experimental conditions in which neither of these methods could have been applied. Specifically, we imaged living *C. elegans* nematodes freely moving in an agarose gel plate, an essay requiring real-time focus correction (which precludes the use of image contrast methods) and containing no fiducial reflective plane (which excludes the use of triangulation approaches). When using RAPID, the worm was kept in focus in the entire area of the gel plate, even at velocities of approximately 400 µm/s, in contrast to images obtained without constant focus monitoring (Figs. 4a,b, and Supplementary Video 4). Furthermore, we were able to use the trace of the objective position required to keep the worm in focus, together with the XY position traces, to determine the 3D trajectory of the nematode (Fig. 4c). From this trajectory, it was possible to analyze the kinematics of the worm motion in three dimensions, for instance correlating the elevation angle over the horizontal plane with the absolute speed. The data show an inverse proportionality between the two, highlighting that worm speed is strongly reduced when climbing the steepest parts of the substrate (Figs. 4d,e).

**Figure 4.**
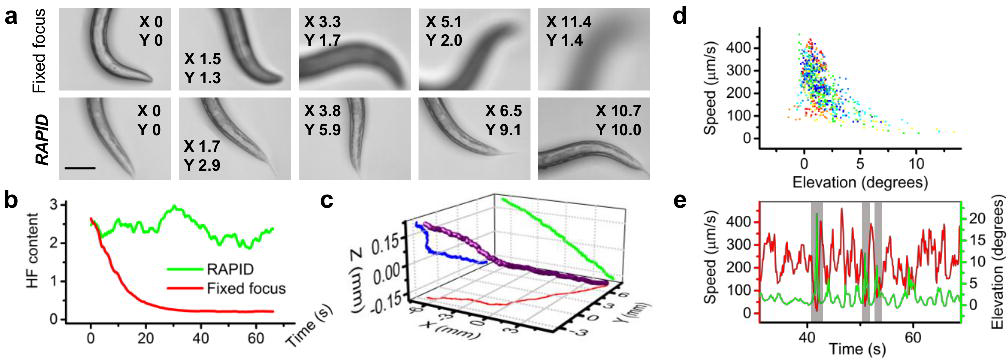
RAPID autofocusing and 3D tracking of fast-moving nematodes. RAPID autofocusing in imaging of fast-moving nematodes (a); the displayed images were acquired at different XY (planar) positions (in mm). High-frequency (HF) content of the images as a function of time (b). RAPID-enabled 3D tracking of the worm (c). A scatter plot of the absolute speed of the worm plotted against the elevation angle (the angle between worm speed direction and the horizontal plane) is shown (d). Dots are colored according to experimental time (blue to red). Exemplar portion of the temporal traces of absolute speed and of elevation angle (e). Gray areas highlight moments of high elevation angle, which corresponds to reduction in absolute speed. Scale bar: 100 µm.

Notably, this three-dimensional analysis of worm motion is beyond the capabilities of existing state-of-the-art AF methods. RAPID offers the possibility to study freely moving microscopic samples in realistic 3D environments, a concept hitherto outside the scope of conventional optical methods.

### RAPID autofocusing in light-sheet microscopy

As a final evaluation, we tested RAPID in high-resolution LSM of cleared mouse brains. In this case, specimen-induced aberrations can introduce a misalignment between the focal plane of the detection objective and the sheet of light (which needs to be coincident), frustrating the very principle of LSM and leading to the acquisition of blurred images. In particular, in tiled imaging of large cleared specimens^12^, defocus can vary between different tiles as well as with tile depth (Supplementary Fig. 6), thus requiring new focus optimization for each tile and at several depths. RAPID could provide focus correction in LSM without adding extra imaging time, maintaining high image quality across the entire specimen (Fig. 5a,b). By using automatic focus correction, we achieved statistically significant contrast enhancement across different tiles (18.5% ± 0.2%, 18000 images) and a contrast increase exceeding 50% in 4% of the images (Fig. 5c). Contrast enhancement was also significant along a single tile (Fig. 5d,e, Supplementary Video 5). In addition, RAPID preserved the image resolution, which can be lost without defocus correction, as demonstrated when imaging samples showing uniform labeling (Fig. 5f). Contrast-based AF has been successfully applied to LSM with comparable results^12,13^, but at the cost of 5% to 50% additional imaging time (Supplementary Note 4).

**Figure 5.**
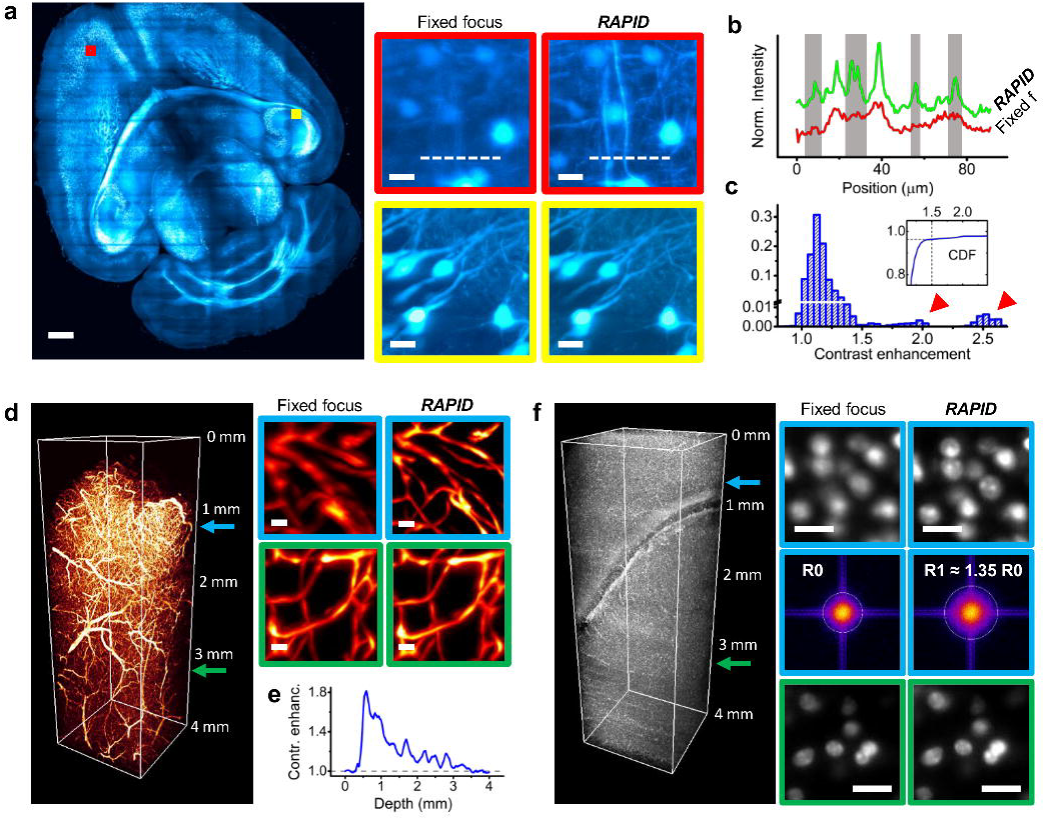
RAPID autofocusing in high-resolution light-sheet microscopy. A virtual slab (500 µm thick) from the brain of a thy1-GFP-M transgenic mouse (a). RAPID defocus correction across different tiles (insets). The intensity profiles were obtained along the dashed lines (b). The gray regions denote fine sample details lost without autofocus. Histogram of contrast enhancement using RAPID relative to fixed focus imaging (c) for all the images forming the slab in (a). The red arrowheads indicate positive outliers, and the inset shows the cumulative density function (CDF). Three-dimensional rendering of an image stack from a vasculature-stained mouse brain showing insets at different depths (d). The RAPID contrast enhancement for this stack as a function of depth is shown in (e). Three-dimensional rendering of an image stack from a mouse brain with nuclear staining. The constant shape of the nuclei allows the evaluation of the resolution enhancement achieved with RAPID by examining the radius of the Fourier transforms (insets, middle line). Scale bars: 1 mm (a), 20 µm (insets).

## Discussion

Albeit automated focus stabilization is ubiquitously used in microscopy, currently used methods are limited in scope, as they are constrained to operate offline^6^ or to monitor a fiducial plane possibly uncorrelated with the sample^7^. Even when both real-time and image-based operation are achieved simultaneously^9,10^, current implementations are still restricted to very specific applications. Here, we reported RAPID, a universal method for real-time image-based AF in optical microcopy. This technique employs the optical principle of phase detection, and can be operated in all wide-field microscopy settings, regardless of the kind of sample, the magnification and the numerical aperture of the system. RAPID is cost-effective, uses only off-the-shelf components and can be easily implemented by users without extensive expertise in optics. The only tuning needed is to find the image registration method that perform best with the collected data. In this respect, the experimenter can choose amongst a wealth of published algorithms and pipelines that have been developed in the last decades^11^, ensuring robust registration outcome also in extreme conditions, e.g. very low signal-to-noise or signal-to-background.

Because RAPID provides real-time readouts of the true focus state of an image, it paves the way for applications beyond the capabilities of conventional methods, e.g., 3D tracking. In this framework, RAPID could also be combined with confocal or two-photon microscopy to maintain moving specimens within the 3D field of view of these techniques without having to image a larger volume^14^.

RAPID also provides a new conceptual and practical framework to measure optical aberrations in wide-field settings. Indeed, the mutual registration of images obtained from two different pupil segments is exploited in this method to determine the lowest significant optical aberration, i.e. defocus. In principle, by splitting the pupil in more regions, the RAPID approach could be used to measure higher-order aberrations in wide-field configuration. Albeit this extension of the method is beyond the scope of this work, we anticipate that it could have a significant impact in the growing field of light-sheet microscopy, allowing real-time correction of specimen-induced aberrations and recovery of diffraction-limited performance^15^.

Finally, being completely agnostic on the illumination strategy and on the sample structure, RAPID can find application in fields other than the life sciences, like material sciences^16^, mineralogy^17^, micro-inspection^18^ or micro-assembly^19^, where automated microscopy is commonly used for high-throughput operation.

## Methods

### RAPID implementation

A 90:10 (transmission:reflection) beam splitter was placed in the infinity-corrected space behind the microscope objective in all experiments except the live imaging of fluorescent yeasts, where a 50:50 beam splitter was used. Light reflected from the beam splitter was sent to a 4f system to create an image of the objective back aperture. The magnification of this 4f system was 200:150 for the BF and EF experiments and 75:200 for the light-sheet experiments. In the secondary pupil plane created by the 4f system, a wedge plate (BSF2550-SIDES-A-SP, Thorlabs, Newton, NJ) was used to spatially separate the two portions of the pupil. A third lens (f = 100 mm for all experiments) was used to create two images of the microscope field of view onto an auxiliary camera, which was Retiga SRV (QImaging, Surrey, BC, Canada) for BF and EF, and Cascade II:512 (Photometrics, Tucson, AZ) for light-sheet experiments. A field stop was placed in the intermediate plane of the 4f system, an image plane of the microscope, to avoid superposition of the two pupil-split images.

The images formed onto two pre-defined portions of the auxiliary camera were mutually aligned by determining the cross-correlation peak. Quality checks of the images and alignment, as well as several image pre-processing strategies were employed to maximize the accuracy and reliability of the system (see Supplementary Methods). The mutual displacement between the pupil-split images was fed to a proportional-integrative feedback loop executed in LabVIEW 2012 (National Instruments, Austin, TX) to correct the objective position. The RAPID software is freely available from https://github.com/ludovicosilvestri/RAPID_CLSM. The hardware and software parameters used in the various experiments presented in this paper are summarized in Supplementary Table 3.

### Histological sample preparation

Samples of atherosclerotic human carotid and human keloid (courtesy of Dr. Cicchi, National Institute of Optics, Italy) were fixed with paraformaldehyde, cut into 5-µm slices with a microtome, stained with standard hematoxylin/eosin, and mounted in glycerol.

### Yeast cultures

The strains used in this study were wild-type *Saccharomyces cerevisiae* (Sigma-Aldrich, St. Louis, MO) and *Schizosaccharomyces pombe* (that express GFP-tubulin under the nmt promoter, courtesy of Prof. Tolić, Ruđer Bošković Institute, Croatia). The yeasts were grown in a standard liquid yeast culture medium (Yeast Peptone D-Glucose) and imaged at 37 °C using a warmed plate. To enhance the expression of GFP, 2 μM of thiamine were added to the growing medium of *Schizosaccharomyces pombe*.

### *C. elegans* motion assay

Wild-type *Caenorhabditis elegans* (C. elegans Behavior Kit, Bio-Rad Laboratories, Hercules, CA) were grown according to the protocol recommended by the supplier. To perform the motion assay, a few *C. elegans* worms were transferred with a spatula onto a fresh agar plate and placed under the microscope. Custom software written in LabVIEW 2012 (available from https://github.com/ludovicosilvestri/RAPID_CLSM) was used to keep the worm in the camera field of view. The same software also recorded the XY positions of the stage and worm in the field of view, providing the absolute XY position of the worm. The Z position was tracked using the position of the Z stage, which was continuously corrected by the RAPID module.

### Mouse brain clearing and staining

Neuronal imaging was performed on cleared brains from thy1-GFP-M transgenic mice (The Jackson Laboratory, Bar Harbor, ME). CLARITY^20^ was used as the clearing procedure. Briefly, the mice were euthanized by overdose of anesthetic (isoflurane) and then transcardially perfused first with 40 ml of 0.01 M of phosphate buffered saline (PBS) solution (pH 7.6) and then with 40 ml of hydrogel solution (4% w/v acrylamide, 0.05% w/v bis-acrylamide, and 0.25% w/v VA044 in PBS). The brains were subsequently extracted and incubated in the same solution at 4 °C for three days. The samples were then degassed in nitrogen atmosphere and incubated at 37 °C to initiate polymerization. The embedded samples were extracted from the gel and incubated in clearing solution (sodium borate buffer 200 mM, sodium dodecyl sulfate 4% w/v, and pH 8.5) at 37 °C with gentle shaking for one month. Before imaging, CLARITY-treated samples were optically cleared using successive incubations in 50 ml of 2,2’-thiodiethanol (30% and 63% v/v) in 0.01 M PBS (TDE/PBS)^21^, each for one day, at 37 °C while gently shaking.

For whole-brain nuclei staining, the CLARITY-processed murine samples were incubated at 37 °C for two days with a 1:50 propidium iodide (P3566, LifeTechnologies, Carlsbad, CA) solution in PBST_0.1_, followed by washing in a PBST_0.1_ solution at 37 °C for one day. Subsequently, they were optically cleared with 63% TDE/PBS before imaging with a light-sheet microscope.

Blood vessels were stained by perfusion with a fluorescent gel used^22^. Mice were euthanized by overdoses of anesthetic (isoflurane) and then transcardially perfused first with 30 ml of a 0.01 M PBS solution (pH 7.6) and then with 60 ml of 4% w/v paraformaldehyde (PFA) in PBS. This was followed by perfusion with 10 ml of a fluorescent gel perfusate containing 0.05% tetramethylrhodamine-conjugated albumin (A23016, Thermo Fisher Scientific, Waltham, MA) as a fluorescent marker. Mice bodies were submerged in ice water, with the heart clamped, to rapidly cool and solidify the gel. Brains were extracted after 30 min of cooling and were incubated overnight in a solution of 4% w/v PFA in PBS at 4 °C. On the next day, brains were rinsed three times with PBS. The fixed brains were incubated in a hydrogel solution for 5 days, followed by degassing and hydrogel polymerization at 37 °C. Subsequently, they were incubated in a clearing solution at 37 °C with gentle shaking for one month. Finally, brains were cleared in 63% TDE for imaging.

All the experimental protocols were designed in accordance with Italian laws and were approved by the Italian Minister of Health (authorization n. 790/2016-PR).

### Bright-field and epifluorescence microscopy

An Eclipse TE300 inverted microscope (Nikon, Tokyo, Japan) equipped with an XYZ stage (L-STEP 13, LANG, Hüttenberg, Germany) was integrated with RAPID for the BF and EF experiments. In the BF modality, a mercury lamp coupled with a red bandpass filter (630/10, Thorlabs) was used to illuminate the sample. In the EF modality, light from a blue LED (M470L3, Thorlabs) was bandpass-filtered (469/35, Semrock, Rochester, NY) to avoid contamination in the fluorescence channel and then reflected to a long-pass dichroic mirror (496 nm edge, Semrock) to illuminate the sample. Light emitted from the sample and transmitted by the dichroic was further bandpass-filtered (520/35, Semrock) to isolate the fluorescence contribution. Images were collected using a sCMOS camera (Orca Flash 2.0, Hamamatsu, Japan). The imaging parameters for the different experiments are summarized in Supplementary Table 3.

### Light-sheet microscopy

The custom-made light-sheet microscope used in the experiments has been described in detail by Müllenbroich and colleagues^23^. Briefly, the sample was illuminated from the side using a virtual light sheet created with a galvo scanner (6220H, Cambridge Technology, Bedford, MA), which was coupled via a 4f system to an air objective (Plan Fluor EPI 10X NA 0.3, Nikon) covered with a protective coverslip. Light emitted from the specimen was detected orthogonally to the illumination plane using an immersion objective corrected for clearing solutions (XLPLN10XSVMP 10X NA 0.6, Olympus, Tokyo, Japan). Then, it was bandpass-filtered to isolate fluorescence light and projected by a tube lens onto the chip of a sCMOS camera (Orca Flash 2.0, Hamamatsu) operating in rolling-shutter mode to guarantee confocal line detection. During imaging, the sample was fixed in a refractive-index-matched quartz cuvette (3/Q/15/TW, Starna Scientific, Hainault, United Kingdom) and moved using a set of high-accuracy linear translators (M-122.2DD, Physik Instrumente, Karlsruhe, Germany). The entire system was controlled by custom software written in LabVIEW 2012 using the Murmex library (Distrio, Amsterdam, The Netherlands). The software can be freely downloaded from https://github.com/ludovicosilvestri/RAPID_CLSM.

### Image analysis

Tiles from whole-slide imaging were stitched together using the FIJI Grid/Collection stitching plugin^24^ (https://fiji.sc). FIJI was also used to produce the images and videos. Three-dimensional rendering was performed with Amira 5.0 (FEI Visualization Sciences Group, Bordeaux, France). The high-frequency content of the nematode time-lapse images was evaluated using a Matlab R2016 script (MathWorks, Natick, MA). Tiled images acquired with LSM were stitched using TeraStitcher^25^ to obtain a low-resolution view of the entire imaging volume. The image contrast of the original images was evaluated using the discrete cosine transform entropy method^13^ and was implemented using Matlab R2016.

## Acknowledgements

We thank Prof. Iva Tolić from Ruđer Bošković Institute (Croatia) for providing the fluorescent *S. Pombe*, and Dr. Riccardo Cicchi from National Institute of Optics (Italy) for providing the histological slides used in this study. This project received funding from the European Union’s H2020 research and innovation programme under grant agreements No. 720270 (Human Brain Project) and 654148 (Laserlab-Europe), and from the EU programme H2020 EXCELLENT SCIENCE - European Research Council (ERC) under grant agreement ID n. 692943 (BrainBIT). The project have also been supported by the Italian Ministry for Education, University, and Research in the framework of the Flagship Project NanoMAX and of Eurobioimaging Italian Nodes (ESFRI research infrastructure), and by “Ente Cassa di Risparmio di Firenze” (private foundation).

## Author contributions

L. Silvestri devised RAPID; L. Silvestri, L. Sacconi and F.S.P. designed the experiments; L. Silvestri and M.C.M. implemented RAPID; I.C. prepared the yeast and nematode samples; A.P.D.G. and I.C. prepared the cleared mouse brains; L. Silvestri, I.C. and A.P.D.G. performed the experiments; L. Silvestri and M.C.M. analyzed the data; L. Silvestri wrote the paper with contributions from all the Authors.

